# HIV-1 Amplifies IL-8 Response of Human Stellate Cells to Gram-Positive Microbial Products Via H4K5 Histone Acetylation

**DOI:** 10.1101/2025.05.12.653460

**Authors:** Lumin Zhang, Rabab O. Ali, Naichaya Chamroonkul, Parissa Tabrizian, Myron Schwartz, Ganesh Gunasekaran, Thomas Schiano, Maria Isabel Fiel, Steven C. Ward, Meena B. Bansal

## Abstract

**Background:** Patients living with human immunodeficiency virus 1 (PLWH) develop accelerated liver fibrosis, but the exact mechanism remains unknown. Activation of hepatic stellate cells (HSCs)—a cornerstone of fibrosis—is influenced by various factors, including viral infection, hepatocellular injury, chronic immune activation, gut barrier dysfunction, and microbial translocation. The role of gram-positive microbial products in human immunodeficiency virus 1 (HIV-1) infection–associated liver inflammation and fibrosis remains poorly understood. This study investigates the effect of lipoteichoic acid (LTA), a major gram-positive bacterial component, on HSCs in the context of HIV-1 infection.

**Methods:** Human HSCs were isolated from liver tissues of both HIV-1–infected and uninfected individuals undergoing hepatic resection. Inflammatory responses of HSCs to LTA stimulation were measured ex vivo via ELISA before and after HIV-1_BaL_ exposure. Western blotting, ChIP-qPCR and RNA-seq were used to reveal the mechanisms contributing to the IL-8 response to LTA stimulation and HIV-1_BaL_ exposure in HSCs.

**Results:** LTA modestly induced interleukin-8/CXCL8 (IL-8) production in HSCs, but this response was significantly heightened following HIV-1 exposure. Increased IL-8 levels were also observed in liver tissues from HIV-1–infected patients. In vitro, IL-8 treatment of HSCs elevated α-SMA and COL1A1 expression, implicating IL-8 in fibrosis progression in HIV-1 infection. Transcriptomic analysis pointed to histone acetylation as a key regulator of the IL-8 response of HSCs to LTA during HIV-1 infection. Supporting this, the histone deacetylase (HDAC) inhibitor Trichostatin A (TSA) further enhanced IL-8 expression in HIV-1–exposed HSCs. ChIP-qPCR confirmed that acetylation of histone H4K5 facilitated IL-8 promoter transactivation, sensitizing HSCs to LTA under HIV-1 influence.

**Conclusions:** HIV-1 infection primes HSCs for an exaggerated response to LTA, driven by histone acetylation and resulting in elevated IL-8 production—potentially accelerating liver fibrosis in PLWH. Given the persistence of microbial translocation despite effective antiretroviral therapy, these findings highlight the need for targeted interventions to prevent or mitigate liver fibrosis in PLWH.

## Introduction

Human immunodeficiency virus 1 (HIV-1) infection is marked by profound depletion of CD4⁺ T cells, leading to systemic immunosuppression. As a result, patients living with HIV-1 (PLWH) are more susceptible to opportunistic and community-acquired bacterial infections. For instance, the risk of *Streptococcus pneumoniae* infection is 100 times higher in HIV-1–infected individuals than in the general population^1^. Early during infection, the loss of gut-associated CD4⁺ T cells compromises intestinal barrier integrity, facilitating the translocation of microbial products into systemic circulation^2^. Notably, this microbial translocation persists even in patients receiving antiretroviral therapy (ART), as ART fails to fully restore gut permeability—making microbial translocation a key driver of sustained immune activation in PLWH.

HIV-induced damage to the gastrointestinal mucosa leads to the translocation of both gram-positive and gram-negative microbial products from the gut lumen into the liver via the portal vein^3^. While gram-negative products typically translocate more efficiently following mucosal disruption^4^, gram-positive bacteria also enter the circulation frequently^5^, either through the same routes or secondary infections. Although extensive research has explored the effects of gram-negative components like lipopolysaccharide (LPS) on immune activation, liver injury, and fibrogenesis in HIV^6,7^, the influence of gram-positive microbial products remains comparatively understudied. Lipoteichoic acid (LTA), a major component of the gram-positive bacterial cell walls, functions as a pathogen-associated molecular pattern (PAMP) capable of modulating host immunity^8–10^. Similar to LPS, LTA triggers CD14-dependent nitric oxide release in vascular smooth muscle cells, macrophages, and mononuclear cells^11^. Elevated serum LTA levels have been detected in PLWH, correlating with heightened autoantibody production and immune activation^12^. Given the liver’s frontline exposure to gut-derived microbial products via the portal vein, the impact of LTA—particularly on hepatic stellate cells (HSCs)—in HIV-associated chronic liver disease remains insufficiently explored.

End-stage liver disease continues to be a major cause of mortality in HIV-positive patients despite successful ART, highlighting ongoing hepatic dysfunction and progressive fibrosis^13^. Numerous studies have shown a positive correlation between circulating HIV RNA levels and advanced fibrosis in HIV mono-infected patients^14–16^. Additionally, co-infection with hepatitis C virus (HCV) accelerates fibrosis progression threefold compared to HCV mono-infection^17^. While cross-talk between HSCs and liver macrophages is critical for HSC activation^18^, HSCs are the primary mediators of hepatic fibrogenesis. Our prior work^19,20^ revealed that HIV-1 infection enhances the sensitivity of liver macrophages to gram-negative microbial stimuli such as LPS, promoting elevated production of pro-inflammatory cytokines including interleukin-8 (IL-8/CXCL8). However, whether these cytokines directly stimulate HSC activation remains unclear.

In this study, we examined the inflammatory responses of primary human HSCs—isolated from liver tissues of both HIV-1–infected and uninfected individuals —upon LTA stimulation. We demonstrate that HIV-1 infection amplifies IL-8 production in response to LTA and that IL-8 exerts pro-fibrogenic effects in HIV-1–infected liver tissue. Furthermore, we identify histone H4K5 acetylation as a mediator of IL-8 transactivation in HIV-1–exposed HSCs, offering mechanistic insight into the epigenetic regulation of liver fibrosis in the context of HIV infection.

## Materials and Methods

### Ethics Statement

De-identified liver resection specimens from HIV-1–positive individuals (nL=L7) with effectively suppressed viral loads on ART were provided by Drs. Schwartz, Ganeskaran, and Fiel. Control liver tissues (nL=L10) were obtained from the Mount Sinai Biobank and included samples from patients undergoing liver resection for colon cancer metastases (nL=L9) or cholangiocarcinoma (n=1). All tissue samples were collected during liver transplantation or tumor resection procedures and were processed as formalin-fixed, frozen, or fresh in Dulbecco’s Modified Eagle Medium (DMEM). The study was approved by the Mount Sinai Institutional Review Board (GCO #06-0523 and #10-1211). Liver tissue samples from both HIV-1–infected and uninfected individuals were used for IL-8 immunohistochemistry (IHC). In addition, blood samples were collected from HIV-positive participants enrolled in a metabolic dysfunction-associated steatotic liver disease (MASLD) study. Population characteristics (Table 2) were analyzed using the R packages *survey*^21^ and *tableone*^22^.

**Table 1:**
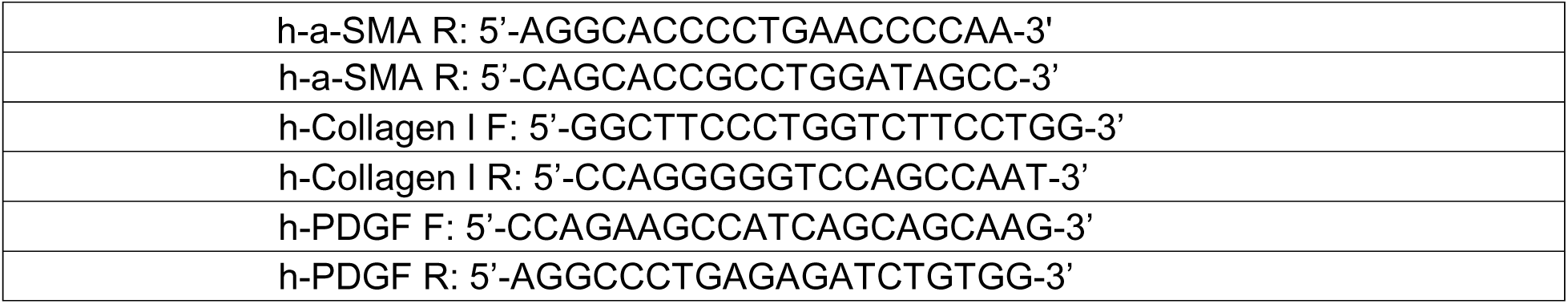
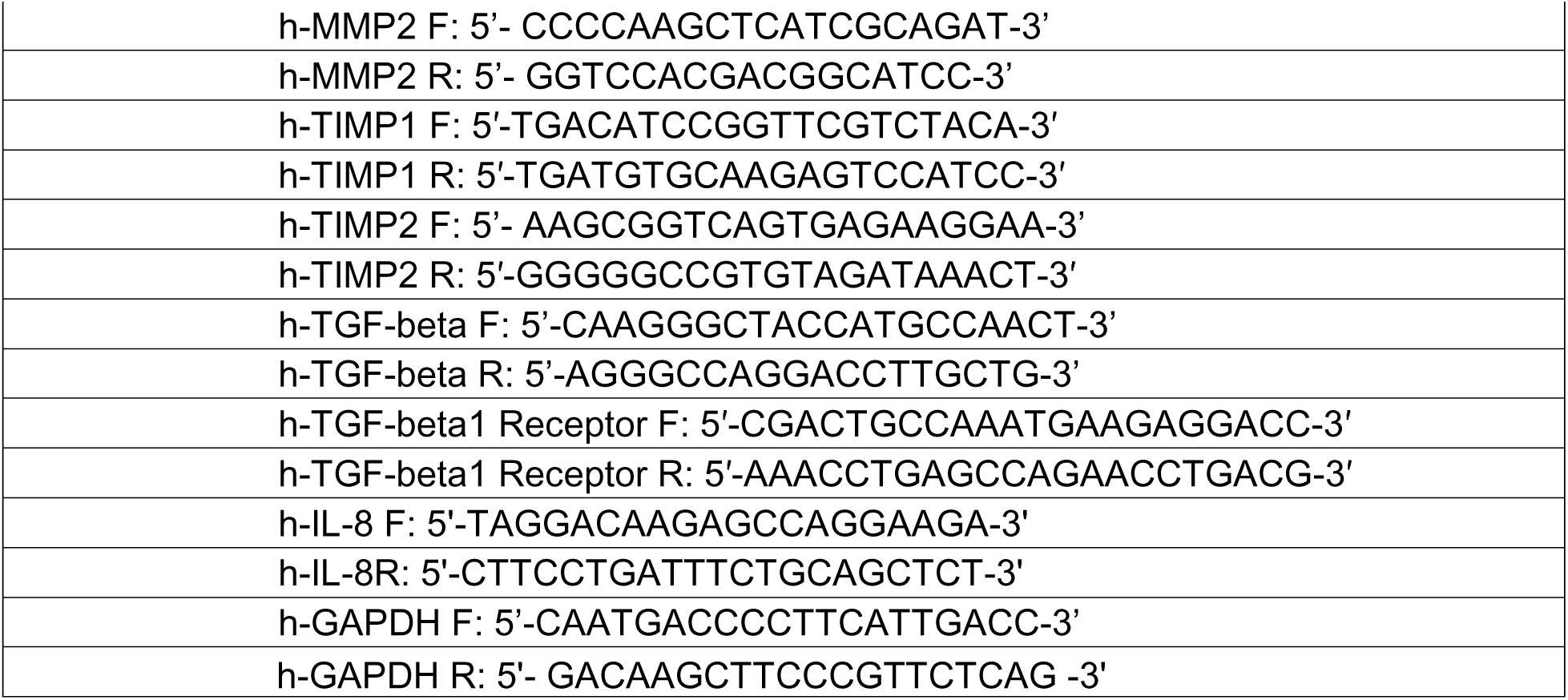
Primers: All primer sequences.

**Table 2:**
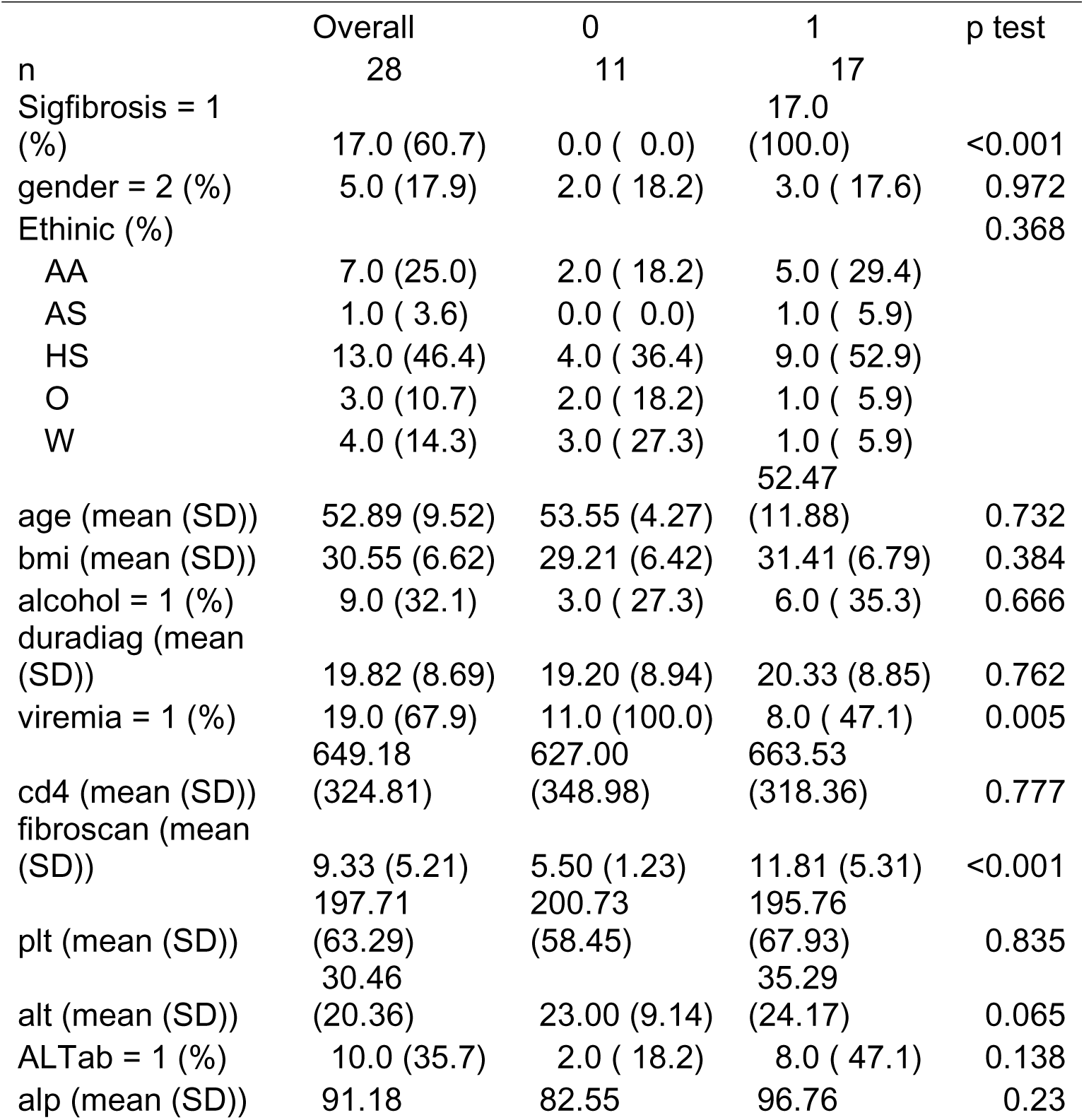
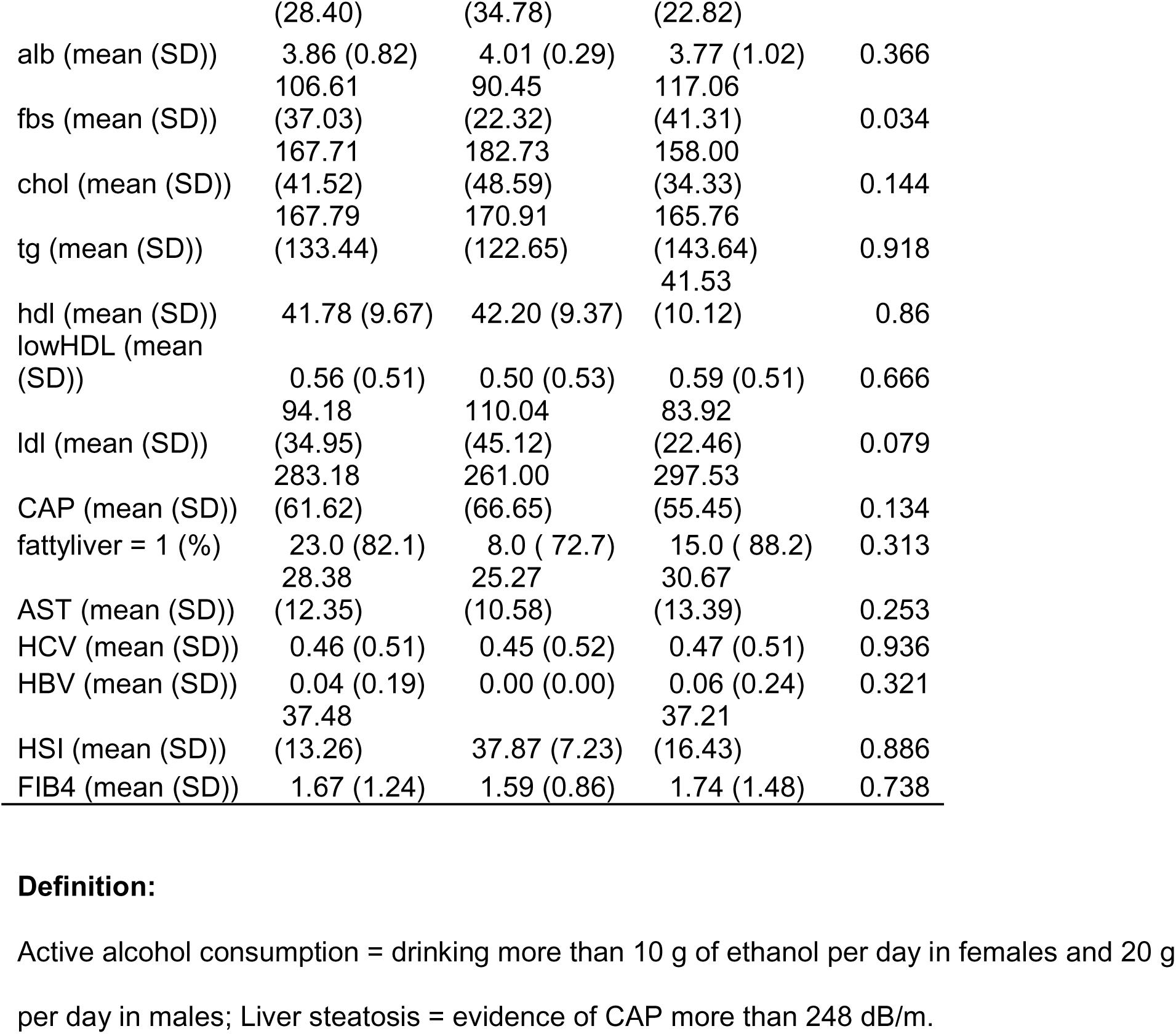
Characteristics of blood donors.

### Isolation of Primary Human Hepatic Stellate Cells

Primary HSCs (pHSCs) were isolated from surgical liver tissue specimens using collagenase perfusion and tissue mincing. Cells were washed with PBS (4×) and centrifuged at 300L×Lg for 5 minutes. Following resuspension in DMEM, cells underwent density gradient centrifugation with 25% and 50% Percoll (Pharmacia, Sweden) as described previously^23^. pHSCs were enriched by adherence to tissue culture plates and cultured for two weeks. Purity was confirmed via α-SMA immunostaining.

### Tissue Immunohistochemistry

Liver tissue sections (prepared in duplicate) were rehydrated, processed for antigen retrieval, and incubated overnight at 4°C with anti–human IL-8 antibody (NOVUS, Cat. NBP2-33819, 1:200 dilution) in a humidified chamber. IL-8 staining was detected using the ImmPress™ Duet Double Staining Polymer Kit (Cat. MP-7724). Four random fields per section were imaged at 100× magnification using Zen software with an LSM 880 laser-scanning microscope. IL-8 signal intensity was quantified using Fiji ImageJ (NIH, Bethesda, MD).

### Flow Cytometry Analysis

Expression of CXCR1, CXCR2, and TLR2 was assessed in pHSCs before and after gp120 stimulation (100ng/mL). Cells (0.5–1×10/mL) were resuspended in PBS with 0.5% bovine serum albumin (BSA), blocked with FcR reagent, and stained with anti-human CXCR1-APC (Biolegend, Cat. 320612), CXCR2-FITC (Biolegend, Cat. 320704), and TLR2-PE (Biolegend, Cat. 309708) for 15Lminutes. After fixation and permeabilization (20 minutes at 4°C), samples were analyzed on an LSRII flow cytometer (BD Biosciences). Data were processed using FCS Express 6 (De Novo Software).

### Histone Extraction and Acetylation ELISA

pHSCs and Lx2 cells (1×10) were seeded in 10 cm dishes and infected with HIV-1_BaL_ virus (NIH AIDS Reagent program, Bethesda, MD) for 48 hours. Total histones (200 ng) were extracted using the EpiQuick™ Total Histone Extraction Kit (Cat. OP-0006) and quantified for acetylation using a histone acetylation ELISA Kit (Cat. P-4022-96) per manufacturer protocols.

### RNA Extraction and RT-qPCR

Total RNA was extracted using RNeasy spin columns (Qiagen, Cat. 74136) and treated with RNase-free DNase (Qiagen, Chatsworth, CA). Reverse transcription was performed using the Clontech RNA-to-cDNA EcoDry Kit. RT-qPCR was conducted on a Roche LightCycler 480 II using SYBR Green PCR master mix (Bio-Rad, Cat. 1708884) with primers at 300 nM (primer sequences in Table 1). Reactions were run in triplicate. Cycle thresholds (CTs) were normalized to α-actin.

### IL-8 ELISA

HSCs were treated under various conditions for 24 hours. Supernatants were collected at designated time points, and IL-8 levels were measured using the Human IL-8 ELISA Kit (Affymetrix, Cat. 88-8086-88).

### CHIP RT-PCR Assay

pHSCs (1×10) were infected with HIV-1_BaL_ virus (NIH AIDS Reagent program, Bethesda, MD) for 48 hours. Chromatin was prepared using the EZ-Zyme™ Chromatin Prep Kit (Cat. 17-375), and ChIP assays were performed with Magna ChIP™ G Kits (Cat. 17409). IL-8 promoter activation was quantified using SYBR Green RT-PCR. CHIP-seq data was analyzed with *Bowtie*2^24^, *Macs callpeak*^25^, *Samtools*^26^, *Bedtools*^27^, and *deepTools*^28^. H4K5 peaks were visualized using IGV.

### RNA-Seq Analysis

Total RNA from Lx2 cells treated with medium, HIV-1_BaL_ (MOI=0.1), and/or LTA (1μg/mL) was sequenced (n=3 per group). RNA quality was assessed by a Nano Drop 2000 Bioanalyzer, and RNA-Seq was performed by Mount Sinai NGS Core. Reads were aligned with STAR^29^, quantified with HTSeq^30^, and analyzed with DESeq2^31^ using hg19 annotation. Pathway analysis used Cytoscape with ClueGO 14 and R packages: *clusterProfiler*^32^, *enrichplot*^33^, and *Weighted Gene Co-Expression Network Analysis* (WGCNA)^34^.

### Western Blot Analysis

Proteins from liver tissues were extracted with T-PER Reagent (Thermo, Cat. 78510). Equal protein amounts were loaded onto NuPAGE 4–12% Bis-Tris gels, transferred to PVDF membranes. Membranes were washed with blotting buffer (1× 0.1% Tween20 PBS) and blocked for 60 minutes in blotting buffer containing 10% low-fat powdered milk. Membranes were washed 3 times with blotting buffer. Blots were probed with primary antibodies (1:1000) overnight at 4°C and secondary antibodies (1:1000) for 1 hour at room temperature. Target protein signals were developed with TMB (SeraCare Life Sciences Inc., Milford, MA) and visualized using the Amersham Imager 600. Antibodies included α-SMA (NOVUS, Cat. NBP 1-45433 and NB100-56565), Collagen I (NOVUS, Cat. NBP 2-12446), MMP9 (NOVUS), and GAPDH (Santa Cruz, Cat. sc-66163). Densitometry was performed using ImageJ and normalized to β-actin.

## Statistical Analysis

Data are reported as meanL±LSEM. Statistical analyses were performed using GraphPad Prism 5.0 (La Jolla, CA). Two-group comparisons used Student’s t test or the Wilcoxon signed-rank test, as appropriate. Multi-group comparisons used one-way ANOVA with Kruskal–Wallis test. Co-localization in IHC was quantified using Fiji ImageJ and Pearson’s *r*. PL<L0.05 was considered statistically significant; P=0.05–0.09 was considered a trend. Non-significant differences are noted as P≥0.1 unless stated otherwise.

## Results

### HIV-1_BaL_ Exposure Amplifies IL-8 Response of pHSCs to LTA Stimulation

To examine how lipoteichoic acid (LTA) affects primary human hepatic stellate cells (pHSCs), three independent control pHSC lines (from HIV-1 uninfected liver tissues) were treated with LTA. Secretion of pro-inflammatory cytokines IL-6, IL-8, IL-10, and TNF-α was assessed before and after 48-hour exposure to HIV-1_BaL_. LTA treatment alone did not significantly induce IL-6, TNF-α, or IL-10 secretion in either condition (Supplemental Fig. 1). However, IL-8 secretion was significantly elevated in HIV-1_BaL_–exposed pHSCs following LTA stimulation compared to LTA alone (Fig. 1A).

**Figure 1:**
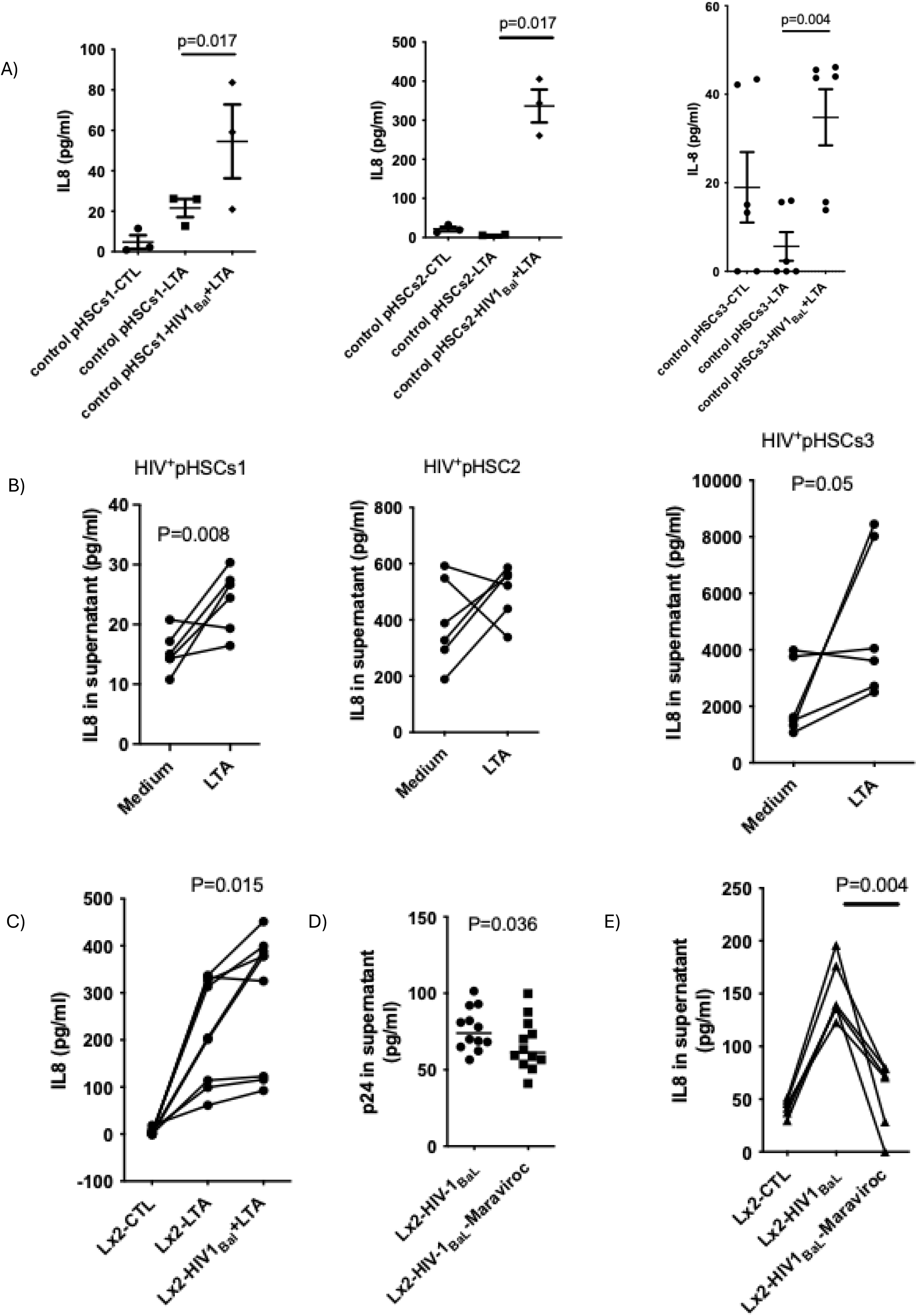
HIV-1 exposure accentuates IL-8 response to LTA stimulation in primary isolated HSCs. A) Primary human hepatic stellate cells (pHSCs) were isolated from HIV-1– uninfected liver tissues (n=3) and treated with LTA (1Lμg/ml) either before or after exposure with HIV-1_BaL_ (MOI=0.1) for 48 hours. Supernatants were collected 24 hours after LTA stimulation, and IL-8 levels were measured by ELISA. B) pHSCs from HIV-1–infected liver tissues (n=6) were stimulated with LTA (1Lμg/ml) for 24 hours, and IL-8 levels were quantified by ELISA. C) Lx2 cells were treated with LTA alone or HIV-1 plus LTA. Supernatants were collected and IL-8 was measured by ELISA (n=6). D) HIV-1 p24 antigen was quantified by ELISA in supernatants from HIV-1_BaL_-exposed Lx2 cells (MOI=0.1) with or without Maraviroc (1LmM) treatment (n=9). E) IL-8 levels were measured by ELISA in HIV-1_BaL_-exposed Lx2 cells treated with or without Maraviroc (1LmM) (n=6). HIV-1_BaL_

To validate these findings, pHSCs isolated from HIV-1–infected livers were stimulated with LTA. Two out of three HIV+ pHSC lines showed significantly increased IL-8 secretion with LTA (Fig. 1B). Similarly, Lx2 cells co-stimulated with LTA and HIV-1_BaL_ displayed synergistic IL-8 induction (Fig. 1C). Lastly, the C-C chemokine receptor type 5 (CCR5) antagonist Maraviroc successfully suppressed HIV-1_BaL_ infection in Lx2 cells, as indicated by reduced p24 levels (Fig. 1D), and significantly attenuated IL-8 production in response to LTA (Fig. 1E).

Flow cytometry analysis of control pHSCs revealed that among IL-8/C-X-C motif chemokine receptors, CXCR1 expression increased upon gp120 stimulation— an HIV-1 envelope protein— while CXCR2 and LTA ligand receptor (TLR2) remained unchanged (Fig. 2A). Notably, neither IL-8 receptor nor TLR2 inhibition altered the heightened IL-8 response to LTA in HIV-1_BaL_– exposed cells (Figs. 2B–2C). Collectively, these findings suggest that HIV-1 sensitizes pHSCs to LTA promoting IL-8 secretion via a pathway independent of IL-8 receptor or TLR2 signaling.

**Figure 2:**
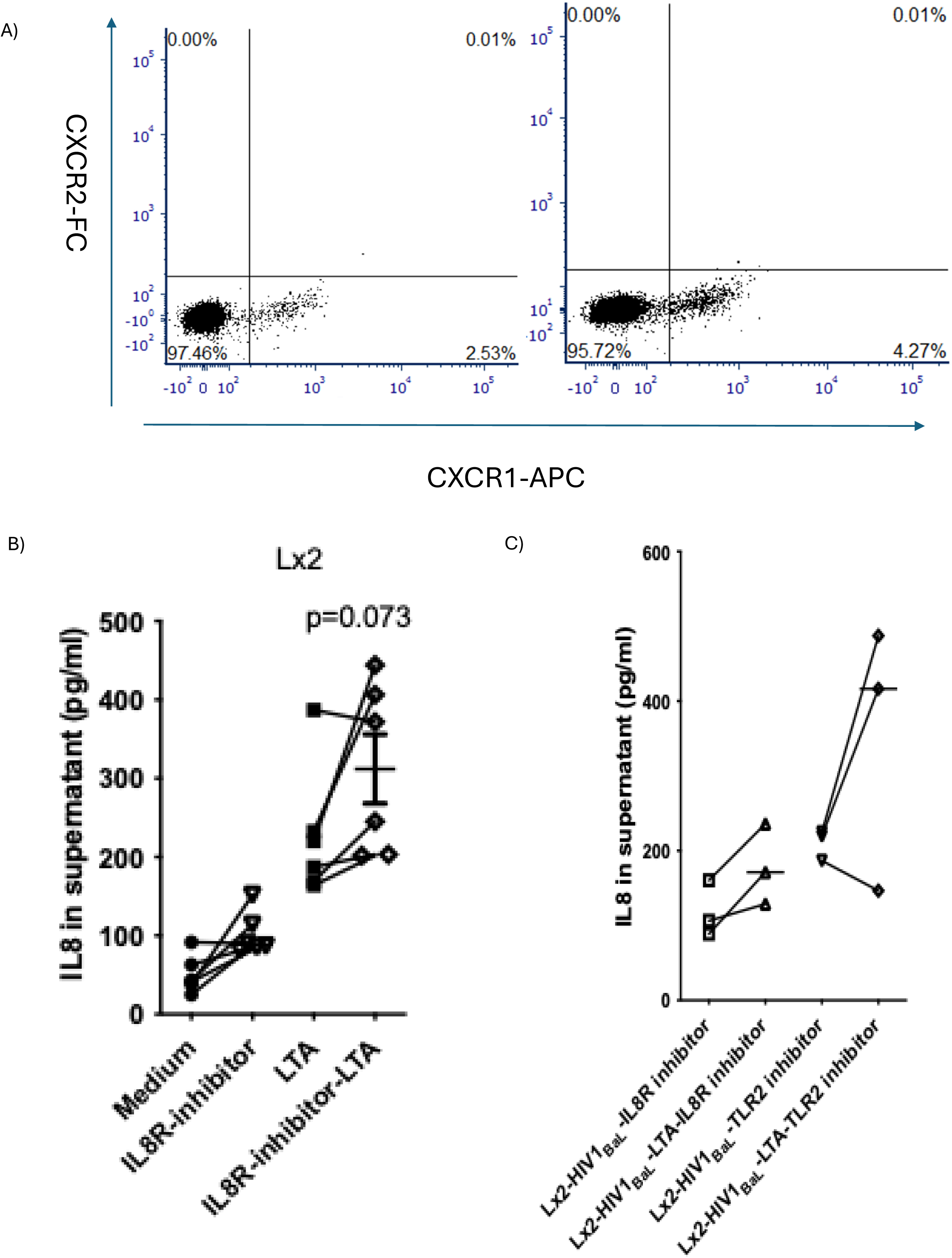
IL-8 response of HSCs to LTA is independent of IL-8 and TLR receptor pathways. A) IL-8 receptor was analyzed by Flow cytometry before and after gp120 stimulation (100ng/ml). B) Lx2 cells were stimulated with LTA (1Lμg/ml) for 24 hours with or without IL-8 receptor inhibitor. Supernatants were analyzed for IL-8 levels by ELISA (n=3). C) Lx2 cells were exposed to HIV-1_BaL_ for 48 hours and then treated with either an IL-8 receptor inhibitor or a TLR2 inhibitor. Supernatants were analyzed for IL-8 levels by ELISA (n=3).

### Elevated IL-8 Expression in HIV-1–Infected Liver Tissue

To confirm the in vitro observations, liver tissues from PLWH were analyzed. IL-8, TGF-β, and α-SMA mRNA levels were significantly higher in HIV-1–infected tissues compared to uninfected controls (Fig. 3A). α-SMA protein levels were also elevated in HIV-1–infected tissues (Fig. 3B). Immunohistochemistry confirmed higher hepatic IL-8 expression in HIV-1–infected tissues (Fig. 3C).

**Figure 3:**
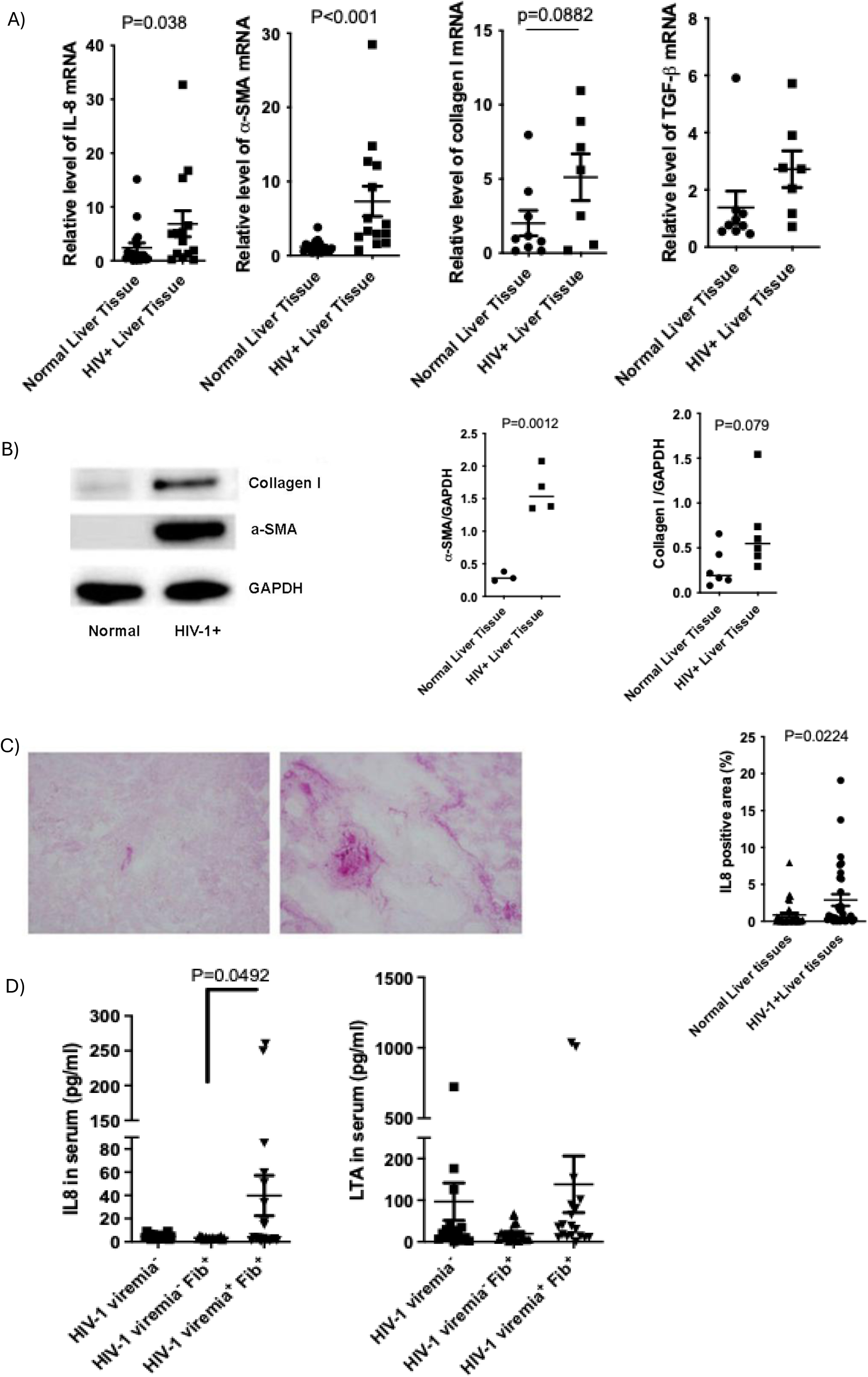
Hepatic IL-8 and pro-fibrogenic gene expression is elevated in HIV-1–infected liver tissues. A) Total RNA was extracted from HIV-1–uninfected (n=7) and HIV-1–infected (n=7) liver tissues. mRNA expression of IL-8, α-SMA, collagen I, and TGF-β was quantified by RT-PCR. B) Protein expression of α-SMA and collagen I was examined by western blot; GAPDH served as a loading control. Band intensity was quantified using ImageJ (n>3). C) IL-8 expression was evaluated by immunohistochemistry in HIV-1–uninfected and HIV-1–infected liver tissues (n=32). The results were quantified by image J. D) Serum IL-8 and LTA concentration were measured in duplicate by ELISA and compared across groups stratified by HIV-1 viremia and fibrosis (n^HIV-1 viremia-^ = 9, n^HIV-1 viremia-Fib+^ = 9, n^HIV-1 viremia+Fib+^ = 10).

Serum IL-8 levels were elevated in individuals with HIV viremia and liver fibrosis compared to those with fibrosis alone. Although mean serum LTA levels showed no groupwise significance, individuals with both HIV viremia and fibrosis exhibited the highest LTA concentrations (mean: 138.81 pg/mL), compared to those with fibrosis only (96.44 pg/mL) or no fibrosis/viremia (19.28 pg/mL) (Fig. 3D).

To test whether LTA or IL-8 exert direct fibrogenic effects, Lx2 cells were treated with increasing doses of LTA and IL-8. IL-8 mRNA expression increased dose-dependently following LTA exposure in Lx2 cells, with no change in other pro-fibrogenic genes (Fig. 4A). IL-8 treatment elevated α-SMA and Collagen I protein levels in Lx2 cells (Fig. 4B), supporting its fibrogenic potential in HIV-associated liver disease.

**Figure 4:**
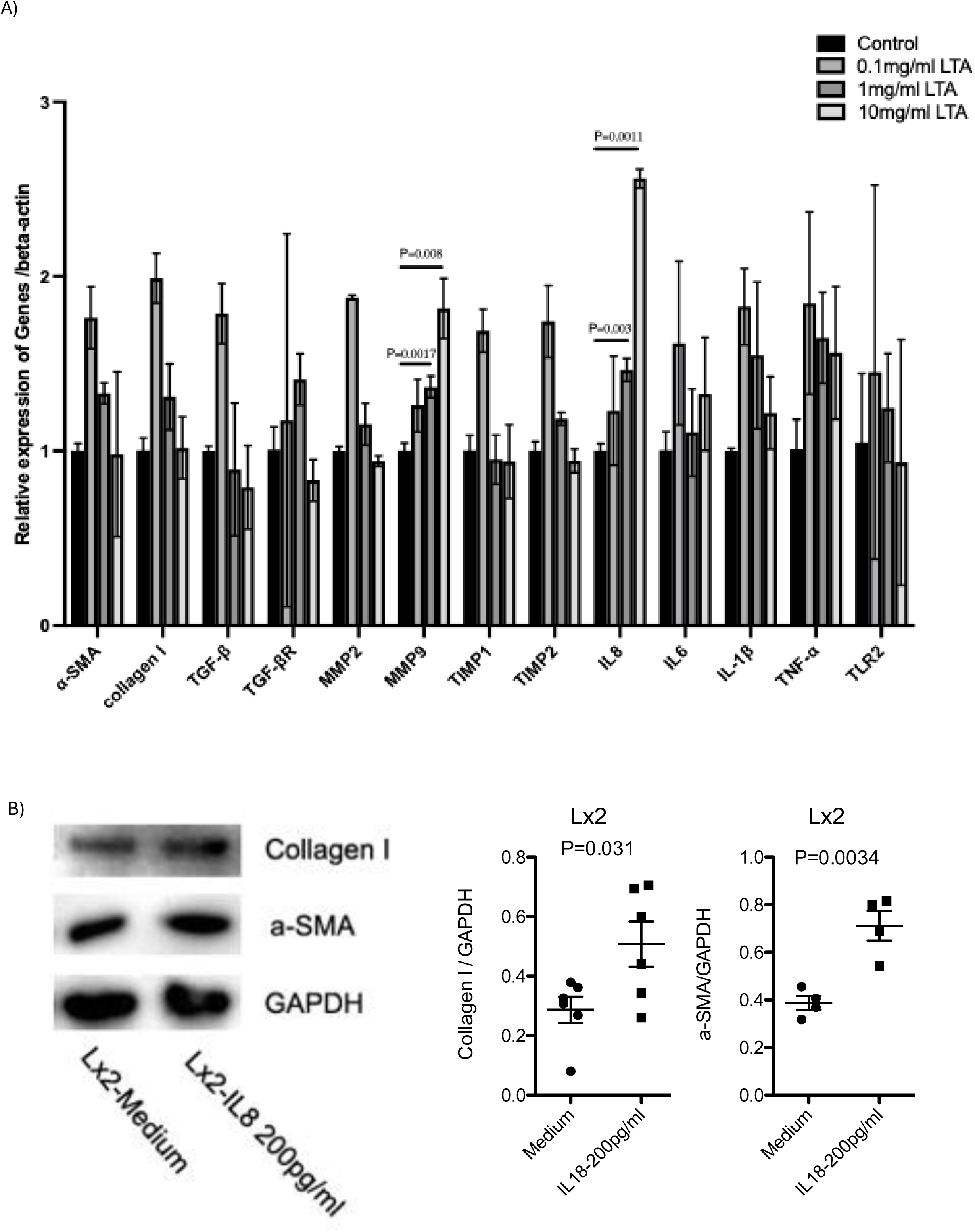
IL-8 increases pro-fibrogenic gene expression in hepatic stellate cell lines. A) Lx2 cells were treated with increasing concentrations of LTA for 24 hours. mRNA expression of pro-fibrotic and inflammatory genes was quantified by SYBR Green RT-PCR. B) Lx2 cells were treated with IL-8 (200Lpg/ml) or PBS x 3. Protein expression of α-SMA and collagen I was analyzed by western blot and normalized to GAPDH. Band intensities were quantified using ImageJ (n=3).

### RNA-Seq Reveals Epigenetic Regulation of IL-8 in HIV-1 and LTA Co-Stimulation

To investigate how HIV-1 exposure sensitizes HSCs to LTA–induced IL-8 expression, RNA-seq was conducted on Lx2 cells under three experimental conditions: untreated, treated with HIV-1_BaL_ alone, or co-treated with HIV-1_BaL_ and LTA (Fig. 5). Although global differential gene expression did not reach statistical significance (Figs. 5A–5B), IL-8 expression trended higher in cells co-stimulated with HIV-1_BaL_ and LTA (p = 0.059) (Fig. 5C).

**Figure 5:**
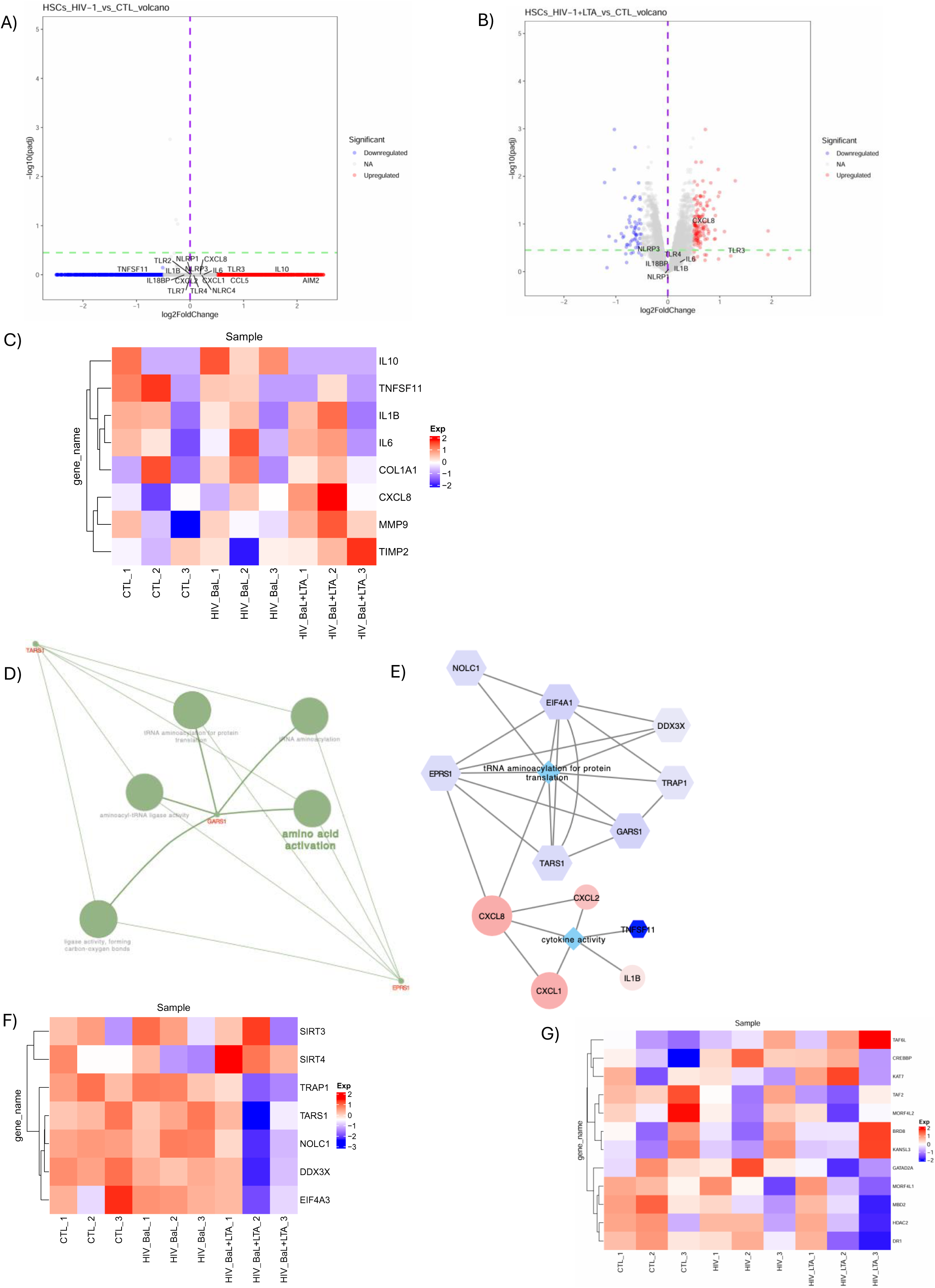
RNA-seq suggests a link between histone acetylation and IL-8 activation following HIV-1 and LTA co-stimulation in Lx2 Cells. A-B) Volcano plots showing differentially expressed genes (DEGs) between control, HIV-1_BaL_-exposed, and HIV-1_BaL_ plus LTA-treated Lx2 cells (pL<L0.05, log2FCL>L1.0). Blue and red dots represent up-and down-regulated genes, respectively. C) Heatmap analysis of pro-inflammatory genes across treatment groups. D) ClueGO pathway enrichment of genes positively correlated with IL-8 expression (correlation coefficient *r*> 0.2), highlighting the tRNA aminoacylation pathway. E) Cytoscape-based network reconstruction of the tRNA aminoacylation pathway, showing IL-8/CXCL8 enrichment. F) Heatmap of tRNA aminoacylation gene expression across three experimental conditions. G) Heatmap analysis of acetylation related genes across three experimental conditions.

WGCNA followed by ClueGO pathway analysis of differentially expressed genes (DEGs) correlated with IL-8 (correlation coefficient *r*>0.2, p≤0.05) linked IL-8 to genes involved in tRNA aminoacylation (Figs. 5D–5E), including an upregulation of *SIRT4* and downregulation of *SIRT3* and related genes (Fig. 5F). Given the role of SIRT3 and SIRT4 in acetylation regulation during oxidative stress and inflammation^35^, we further explored these findings. This revealed altered expression of genes associated with histone H4 acetylation and deacetylation—such as HDAC2 and MORFL2^36,37^—in response to combined HIV-1_BaL_ and LTA exposure (Fig. 5G), suggesting an epigenetic mechanism in IL-8 amplification in LTA and HIV-1 co-stimulated Lx2 cells^35^.

### H4K5 Acetylation is Required for IL-8 Transactivation in HIV-1–Exposed HSCs

To test the role of histone acetylation, the histone deacetylase (HDAC) inhibitor Trichostatin A (TSA) was applied during LTA stimulation. TSA significantly increased IL-8 secretion in both control and HIV+ pHSCs (Figs. 6A–6B) without affecting IL-1β, IL-6, IL-10, or TNF-α (Supplemental Fig. 1). HIV-1_BaL_ infection also elevated total acetylation in Lx2 cells (Supplemental Fig. 2), with H4K5 acetylation specifically increased (Fig. 6C), confirmed by Western blot (Fig. 6D). H3 acetylation remained unchanged (Supplemental Fig. 3A–D).

**Figure 6:**
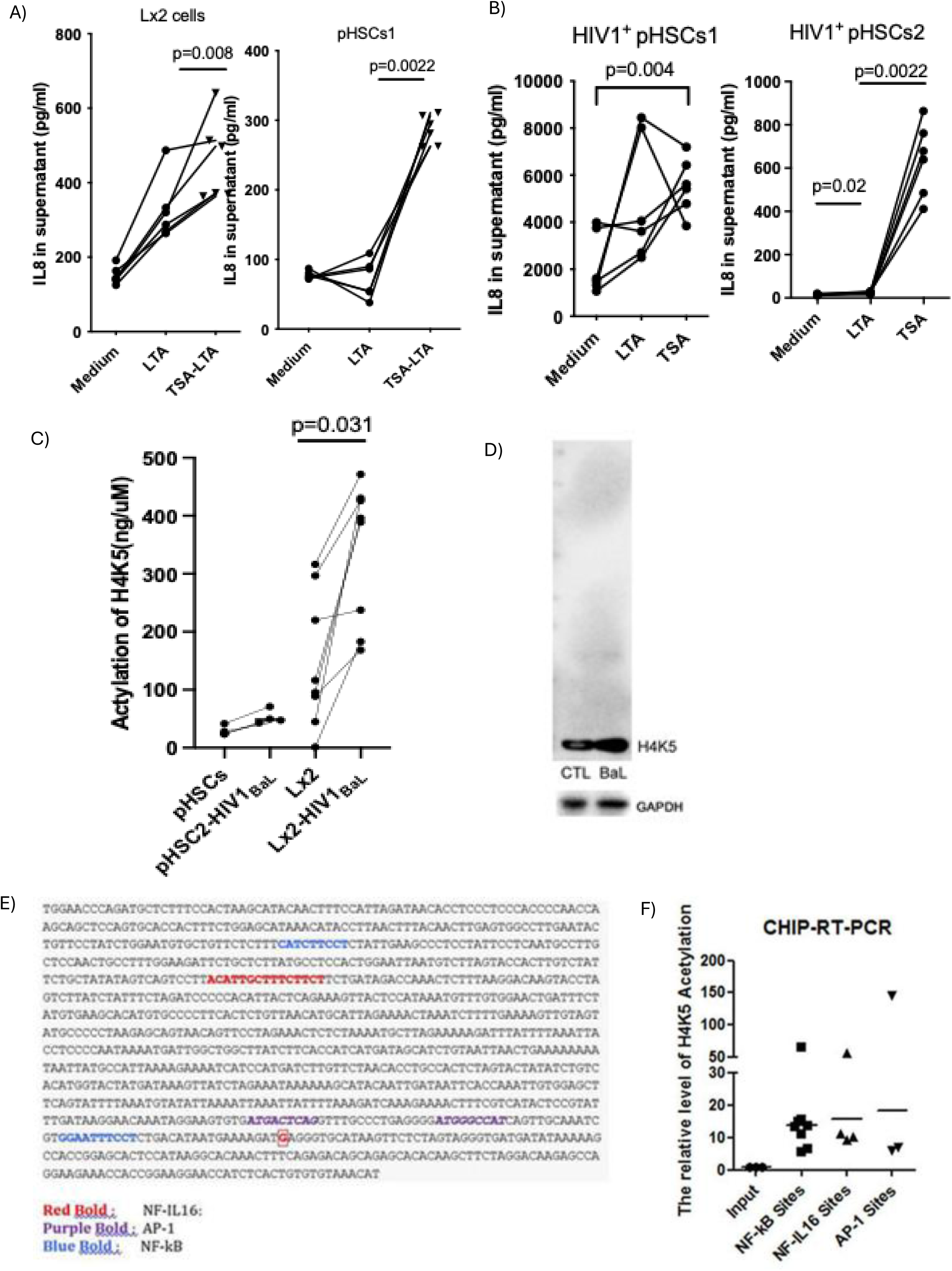
H4K5 acetylation is essential for IL-8 response to LTA stimulation in HIV-1 infected HSCs. A) Two pHSCs isolated from HIV-1–uninfected individuals were stimulated with LTA with or without HDAC inhibitor (TSA) for 24hrs, IL-8 in supernatant was measured by ELISA (n=6). B) Two pHSCs isolated from HIV-1–infected individuals were stimulated with LTA with or without HDAC inhibitor (TSA) for 24hrs, IL-8 in supernatant was measured by ELISA (n=6). C) H4K5 acetylation protein was measured by ELISA in both pHSCs (n=4) and Lx2 cell lines (n=8) before and after HIV-1_BaL_ exposure. D) H4K5 acetylation protein was examined by western blot and normalized to GAPDH. E) Binding motif analysis of IL-8 promoter. F) Chromatin immunoprecipitation (ChIP) was performed using anti-H4K5 antibody. DNA was purified and analyzed by SYBR RT-PCR for enrichment of NF-κB, AP-1, and NF-IL6 binding motifs in the IL-8 promoter.

ChIP-RT-PCR identified H4K5 acetylation enrichment at IL-8 promoter regions in HIV-1– infected Lx2 cells, particularly at NFκB, AP-1, and NF-IL6 binding sites (Figs. 6E–6F). ChIP-seq further confirmed strong peaks in NF-kB and AP-1 binding motifs near the IL-8 transcriptional start sites (Supplemental Fig. 4A–B). Together, these results demonstrate that H4K5 acetylation is a critical mediator of IL-8 transcriptional activation in response to LTA in HIV-1–infected hepatic stellate cells.

## Discussion

End-stage liver disease remains a major cause of mortality among PLWH, with multiple studies showing a positive correlation between HIV viremia and advanced fibrosis in HIV mono-infection^13–16^. Although chronic HIV infection disrupts immune homeostasis and accelerates liver fibrosis, the specific mechanisms linking HIV-mediated inflammation to hepatic fibrogenesis have not been fully elucidated. Our ex vivo and in vitro findings demonstrate that HIV-1 exposure amplifies the IL-8 response of HSCs to LTA—the gram-positive microbial component—with histone acetylation–mediated IL-8 transactivation identified as a key underlying mechanism.

Elevated serum IL-8 levels (≥46.5 pg/mL) have been independently associated with liver-related mortality and the need for liver transplantation^38,39^. Although IL-8 is elevated in mono-infection^40^, its pathological role in HIV-associated liver fibrosis has remained uncertain. Some studies have found no correlation between IL-8 levels and fibrosis in HIV/HCV co-infection, while others suggest elevated IL-8 predicts a greater risk of fibrosis progression^41,42^. In our HIV-1 mono-infected cohort, IL-8 levels were significantly higher in viremic individuals with advanced fibrosis compared to non-viremic counterparts. Interestingly, this distinction was present only in individuals with advanced fibrosis, suggesting a specific link between IL-8 expression and HIV-mediated fibrogenesis. Immunohistochemistry of HIV-1–infected liver tissues confirmed increased hepatic IL-8 expression, which was accompanied by upregulated pro-fibrogenic markers α-SMA and collagen I. In vitro, stellate cell lines responded to IL-8 stimulation with increased expression of these same fibrosis-related genes, supporting a direct fibrogenic role for IL-8 in HSCs.

Previous studies have identified IL-8 as a marker of inflammatory response to liver injury in HSCs pretreated with TNF-α^43,44^. Furthermore, LTA has been shown to promote IL-8 production through NF-κB activation in HSCs pre-primed with TNF-α and IL-1β^45^. Notably, in our study, LTA alone was sufficient to stimulate IL-8 secretion in the absence of such priming, and this effect was significantly enhanced by HIV-1 exposure. Importantly, HIV-1–exposed HSCs did not exhibit increased TNF-α or IL-1β production, reinforcing the possibility that HIV infection independently alters HSC sensitivity to microbial products. Although prior work implicated TLR-mediated interactions between HIV-1, immune cells, and HSCs^46^, our data show that the enhanced IL-8 response is independent of IL-8 receptors or TLR2 signaling. Consistently, RNA-seq analyses revealed no activation of the canonical NF-κB pathway following HIV-1 and LTA co-exposure (Supplementary Fig. 5), suggesting that IL-8 induction involves an alternative regulatory mechanism.

Transcriptomic analysis linked IL-8 transactivation to the aminoacylation pathway and identified differential expression of genes related to histone H4 acetylation. Previous research has shown that IL-8 promoter acetylation contributes to its post-translational regulation^47^and that HDAC inhibitors can suppress IL-8–driven inflammation in various disease contexts^48^. In our ex vivo experiments, treatment with the HDAC inhibitor TSA selectively increased IL-8 responses to LTA, regardless of HIV-1 exposure, without affecting other cytokines such as IL-1β, IL-6, or TNF-α. Through ELISA, ChIP-seq, and ChIP-qPCR, we identified specific enrichment of H4K5 acetylation at the IL-8 promoter in HIV-1–infected cells. This modification promotes chromatin remodeling and recruitment of critical binding sites in the IL-8 promoter i.e., transcription factors AP-1, NF-IL6, and NF-κB^49^.

These findings align with prior studies in cystic fibrosis^50–53^and hepatitis B virus infection^54^, where histone hyperacetylation was shown to drive IL-8 expression in epithelial or hepatocellular cells. Our work highlights the unique role of histone acetylation in modulating HSC responses to microbial stimuli in HIV-associated liver disease. This discovery opens up new avenues for exploring the epigenetic regulation of fibrosis in HIV and underscores the potential therapeutic relevance of targeting gram-positive microbial products or histone acetylation pathways.

## Conclusion

Our ex vivo and in vitro findings underscore the pathological contribution of LTA to hepatic inflammation and fibrogenesis during chronic HIV-1 infection. Elevated serum LTA levels in HIV-1–infected individuals with both viremia and fibrosis, alongside amplified IL-8 secretion from HSCs in response to LTA under HIV exposure, point to gram-positive microbial translocation as a critical driver of disease progression. Transcriptomic analysis revealed that HIV-1 enhances histone H4K5 acetylation, thereby increasing IL-8 promoter activity in response to LTA stimulation, and contributing to sustained fibrogenic signaling in HSCs. These findings link gut-derived microbial products, IL-8–driven inflammation, and epigenetic regulation of HSC activation, providing important mechanistic insight and potential therapeutic targets for managing HIV-associated liver fibrosis.

## Abbreviations

(AST): Aspartate Aminotransferase
(ALT): Alanine Aminotransferase
(ALP): Alkaline Phosphatase
(Alb): Albumin
(ART): Antiretroviral therapy
(BMI): Body Mass Index
(BSA): Bovine serum albumin
(CAP): Controlled Attenuation Parameter
(ChIP-qPCR): Chromatin immunoprecipitation with quantitative PCR
(CT): Cycle threshold
(CCR5): C-C chemokine receptor type 5
(CXCR): C-X-C motif chemokine receptor 1
(DEGs): Differentially expressed genes
(DMEM): Dulbecco’s Modified Eagle Medium
(ELISA): Enzyme-linked immunosorbent assay
(FBS): Fasting Blood Sugar
(FIB-4): Fibrosis-4 Index
(HSC): Hepatic stellate cells
(HBV): Hepatitis B virus
(HCV): Hepatitis C virus
(HIS): Hepatic Steatosis Index
(HDL): High-Density Lipoprotein
(HDAC): Histone deacetylase
(HIV-1): Human immunodeficiency virus 1
(IHC): Immunohistochemistry
(IL-1β): Interleukin-1β
(IL-8): Interleukin-8/CXCL8
(LPS): Lipopolysaccharide
(LTA): Lipoteichoic acid
(LSM): Liver Stiffness Measurement
(LDL): Low-Density Lipoprotein
(MASLD): Metabolic dysfunction-associated steatotic liver disease
(MOI): Multiplicity of infection
(PAMP): Pathogen-associated molecular pattern
(Plt): Platelet count
(pHSCs): Primary HSCs
(PLWH): Patients living with human immunodeficiency virus 1
(RNA-seq): RNA sequencing
(SRT): Sirtuin
(TLR): Toll-Like Receptor
(Chol): Total cholesterol level
(TG): Triglycerides
(TSA): Trichostatin A
(TF): Transcription factor
(TNF-α): Tumor necrosis factor α
(WGCNA): Weighted Gene Co-Expression Network Analysis

## Acknowledgement

We appreciate the reagents support from NIH AIDS Reagents Program.

## Authors’ Contribution

Drs. Zhang and Bansal conceptualized and designed the experiments. Dr. Zhang performed the experiments and conducted bioinformatics analyses. Drs. Zhang, Ali, and Bansal analyzed the data and drafted the manuscript. Dr. Chamroonkul prepared serum samples. Drs. Tabrizian, Schwartz, Gunasekaran, Schiano, and Fiel provided clinical samples and associated clinical data. All authors contributed to the manuscript revisions and approved the final version.

## Data Availability

Processed data files are available at github.com/lumin2003/HIV_LTA_files/.

## Declarations

The human study has been approved by Mount Sinai IRB (GCO #06-0523 and #10-1211).

## Consent for Publication

There is no individual patient information in this study.

