## Supplemental Figures for "HIV-1 Amplifies IL-8 Response of Human Stellate Cells to Gram-Positive Microbial Products Via H4K5 Histone Acetylation" (from another version of this preprint: version 1, 2025-05-15)

### Supplemental Figure 1

A)

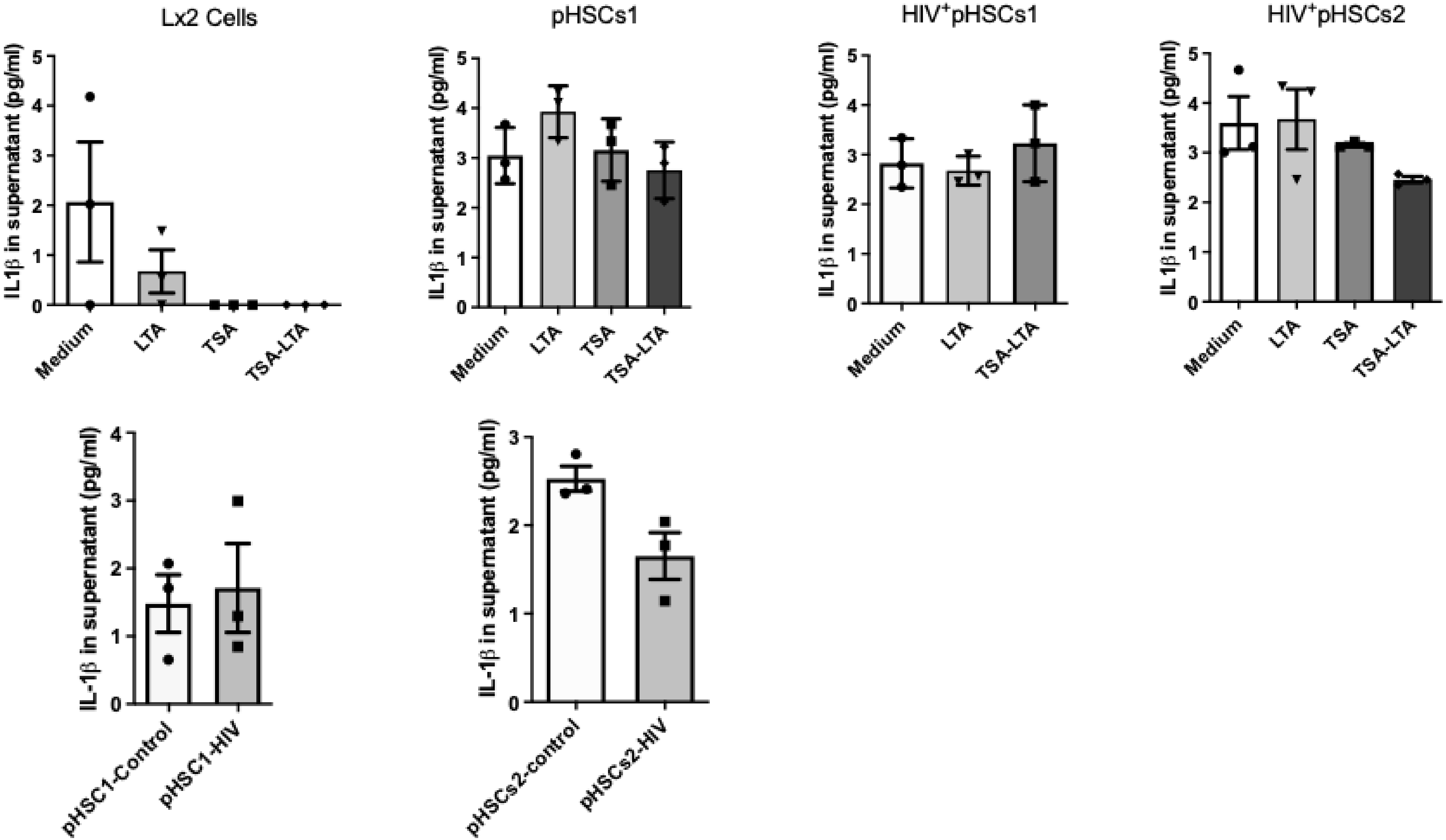

### Supplemental Figure 1

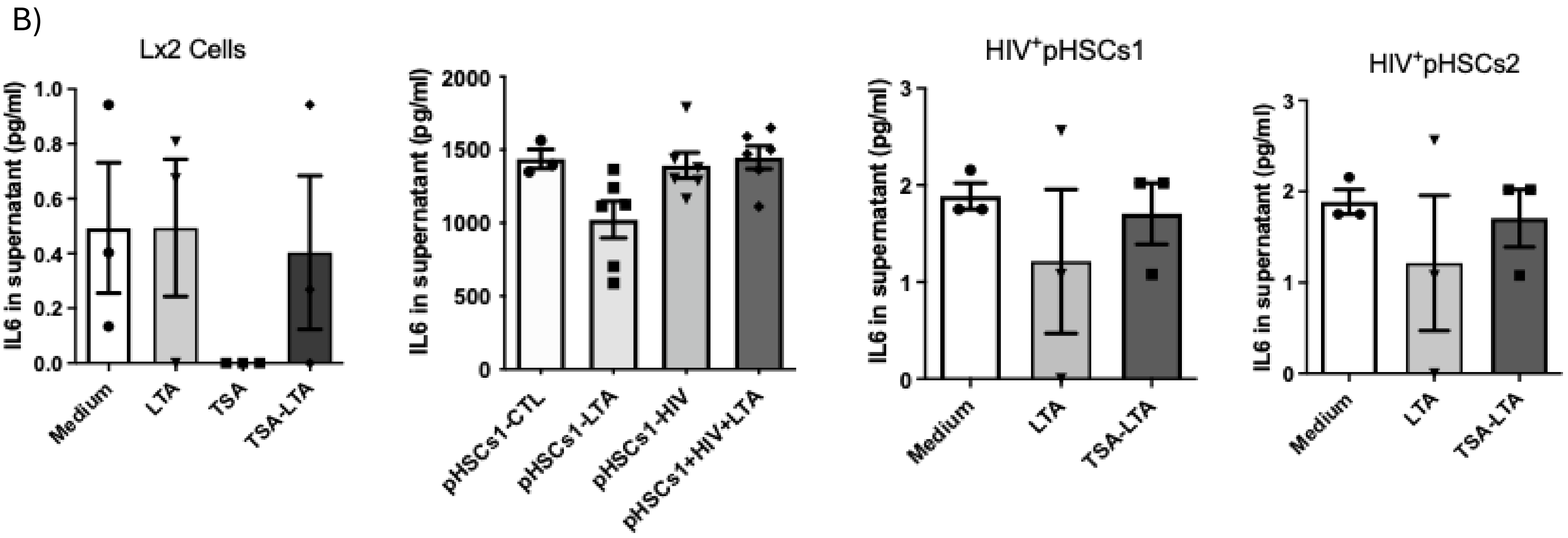


A, B)pHSCs were isolated from HIV-uninfected human liver tissues and treated with LTA (1ug/ml) before and after 2 days of infection with HIV-$1_{BaL}$ (MOI=0.1). Supernatant was collected and used for IL1b (pg/ml) quantification by ELISA (n=3). C) Supernatant was collected at 24-hrs post-LTA stimulation and used for IL6 (pg/ml) by ELISA.

### Supplemental Figure 1

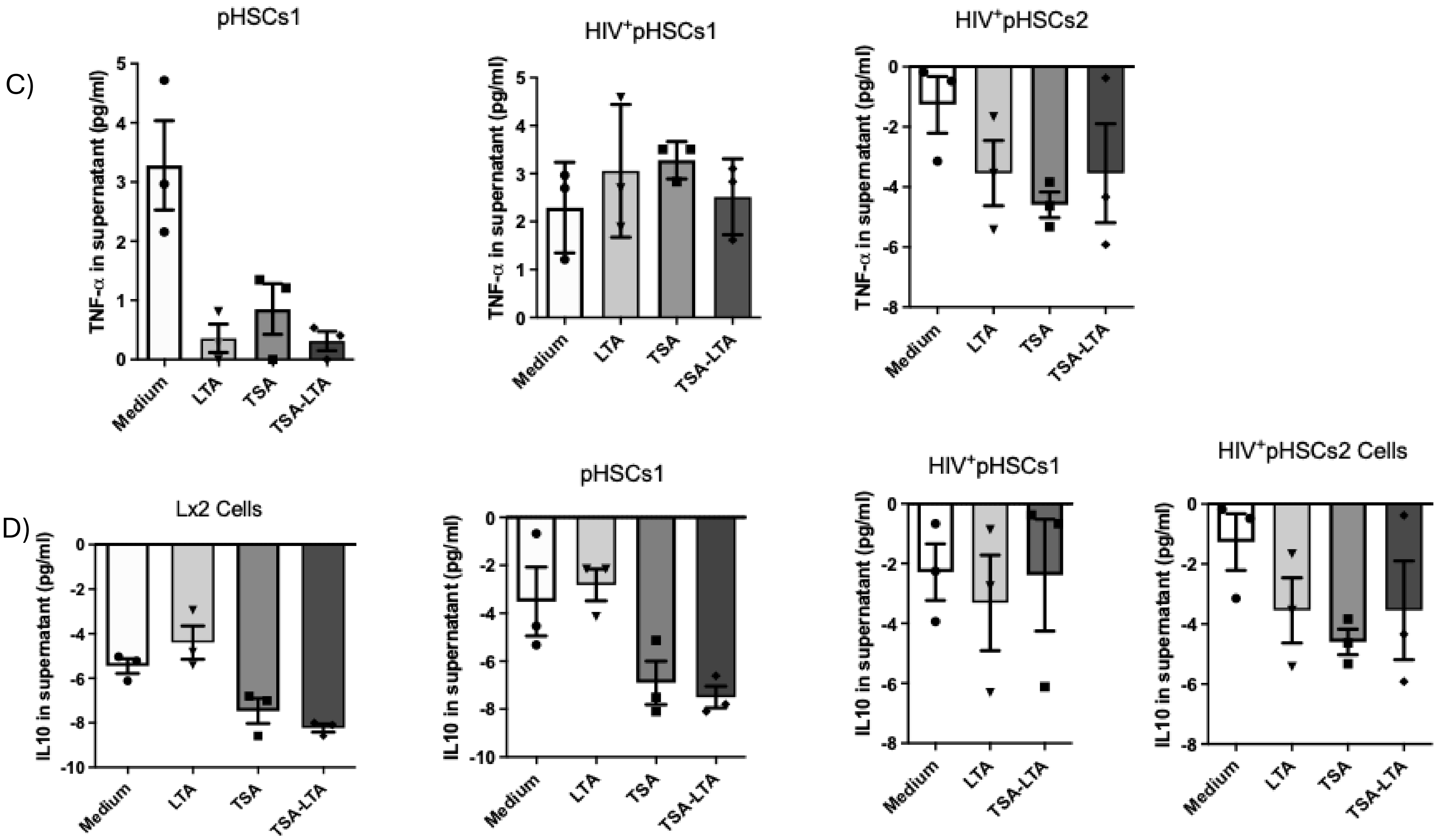

Supernatants were collected from Lx2 cells, primary hepatic stellate cells (pHSCs), and HIV$^+$ pHSCs under the following conditions: untreated, LTA stimulation alone, HIV-1$_{BaL}$ exposure alone, or combined HIV-1$_{BaL}$ exposure and LTA stimulation.

A) IL-1β in supernatant was measured by ELISA (n>=3) ;

B) B) IL-6 in supernatant was measured by ELISA (n>=3);

C) TNF-a in supernatant was measured by ELISA (n>=3) ;

D) IL-10 in supernatant was measured by ELISA (n>=3)

### Supplemental Figure 2

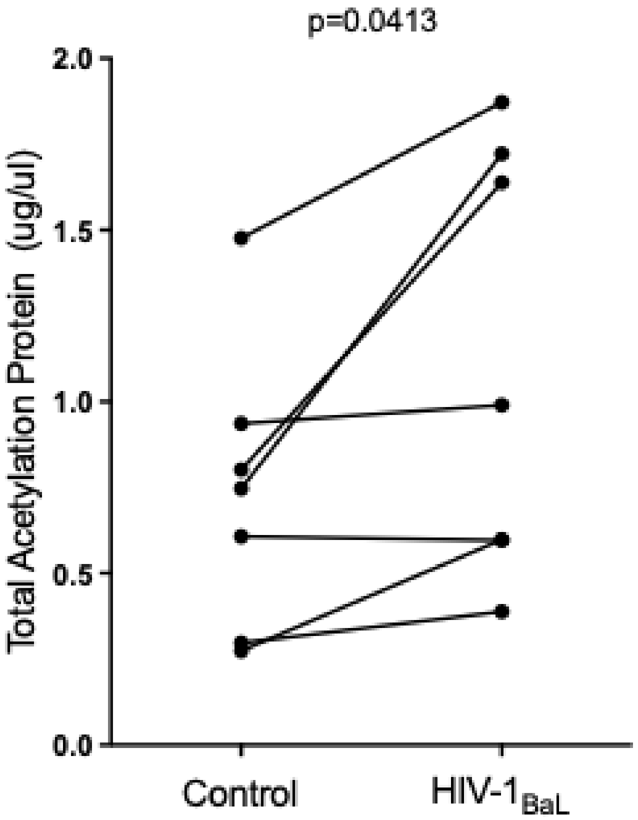


Total acetylation protein was measured by ELISA in both Lx2 cell lines (n=5) before and after HIV-$1_{BaL}$ exposition.

### Supplemental Figure 3

A)

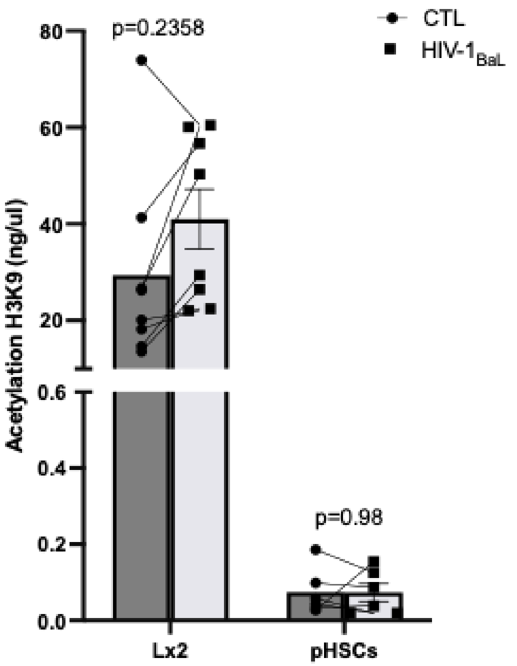


B)

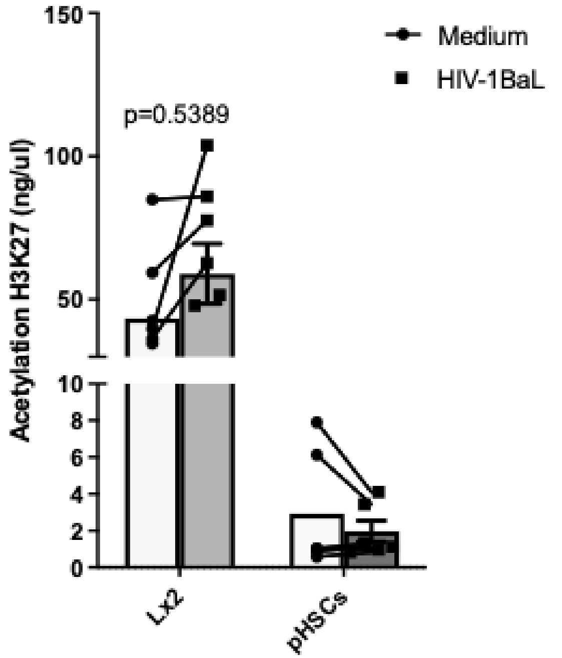


C)

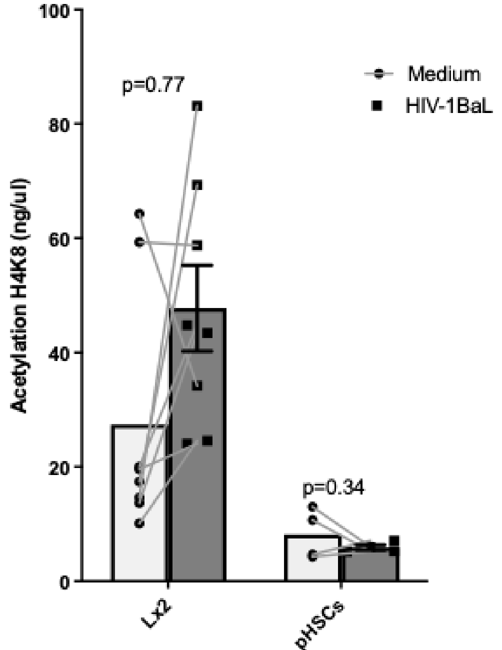


D)

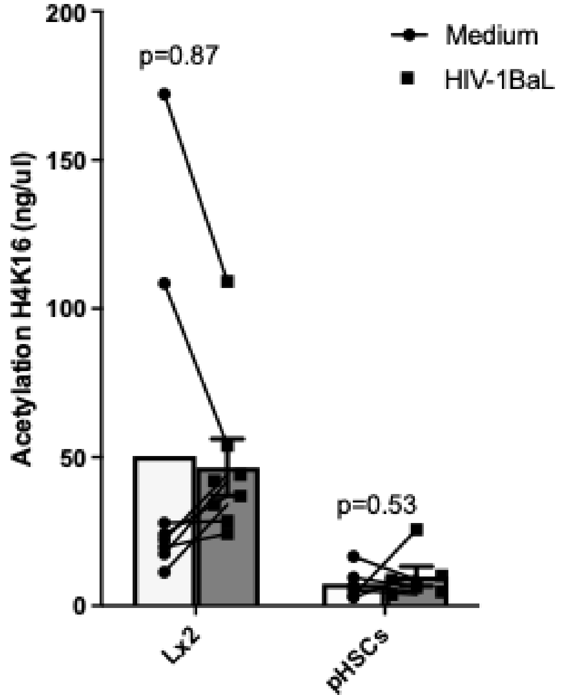

A) H3K9 acetylation protein was measured by ELISA in both pHSCs (n=6) and Lx2 cell lines (n=8) before and after HIV-$1_{BaL}$ exposure.
B) H3K27 acetylation protein was measured by ELISA in both pHSCs (n=6) and Lx2 cell lines (n=8) before and after HIV-$1_{BaL}$ exposure.
C) H4K8 acetylation protein was measured by ELISA in both pHSCs (n=6) and Lx2 cell lines (n=8) before and after HIV-$1_{BaL}$ exposure.
D) H4K16 acetylation protein measured by ELISA in both pHSCs (n=6) and Lx2 cell lines (n=8) before and after HIV-$1_{BaL}$ exposure.

### Supplemental Figure 4

A)

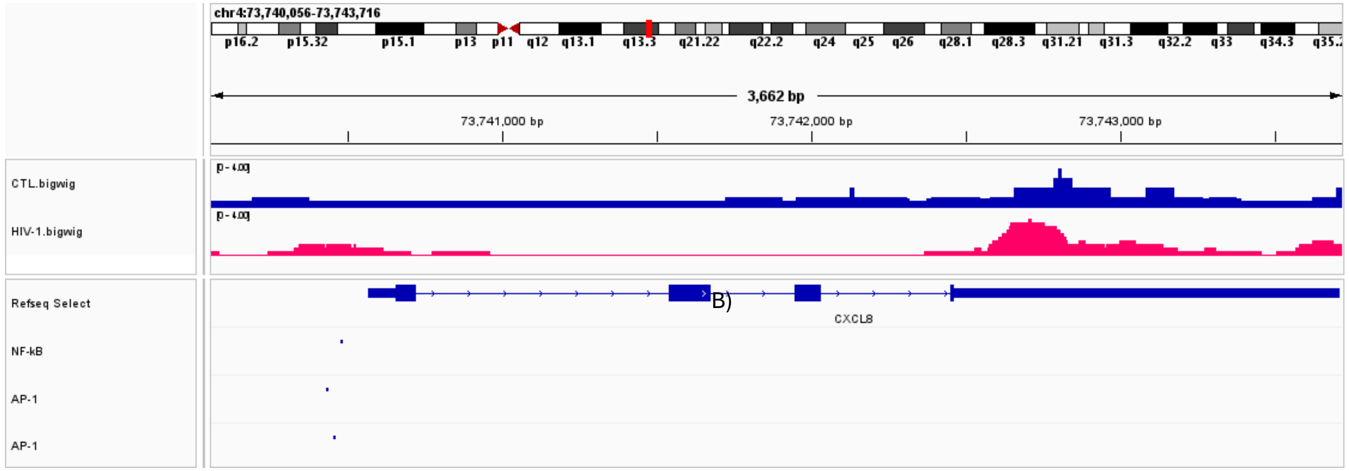


B)

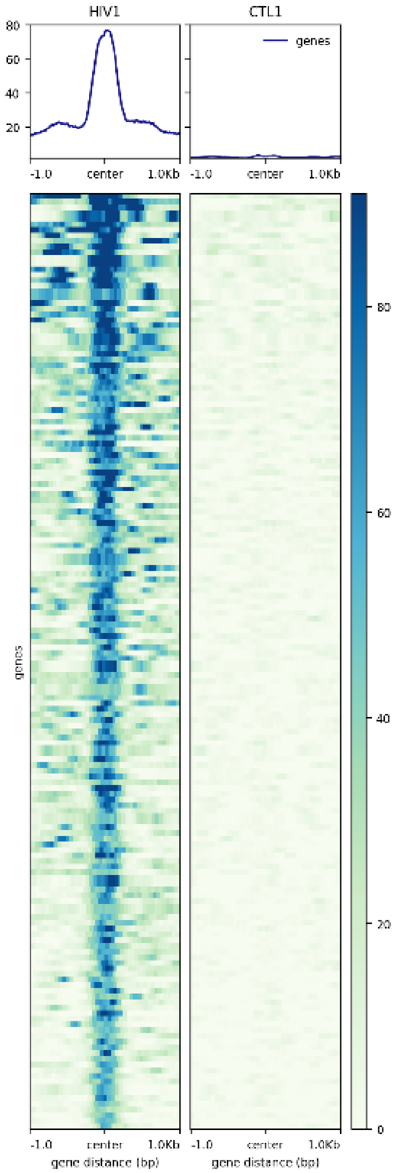


A) Integrative Genomics Viewer (IGV) browser tracks of H4K5 peaks and binding motifs at the CXCL8 locus before and after HIV-1 exposition of Lx2 cells.
B) B) Clustered heatmap of H4K5 enrichment around transcription start sites (±1000 bp) in different conditions.

### Supplemental Figure 5

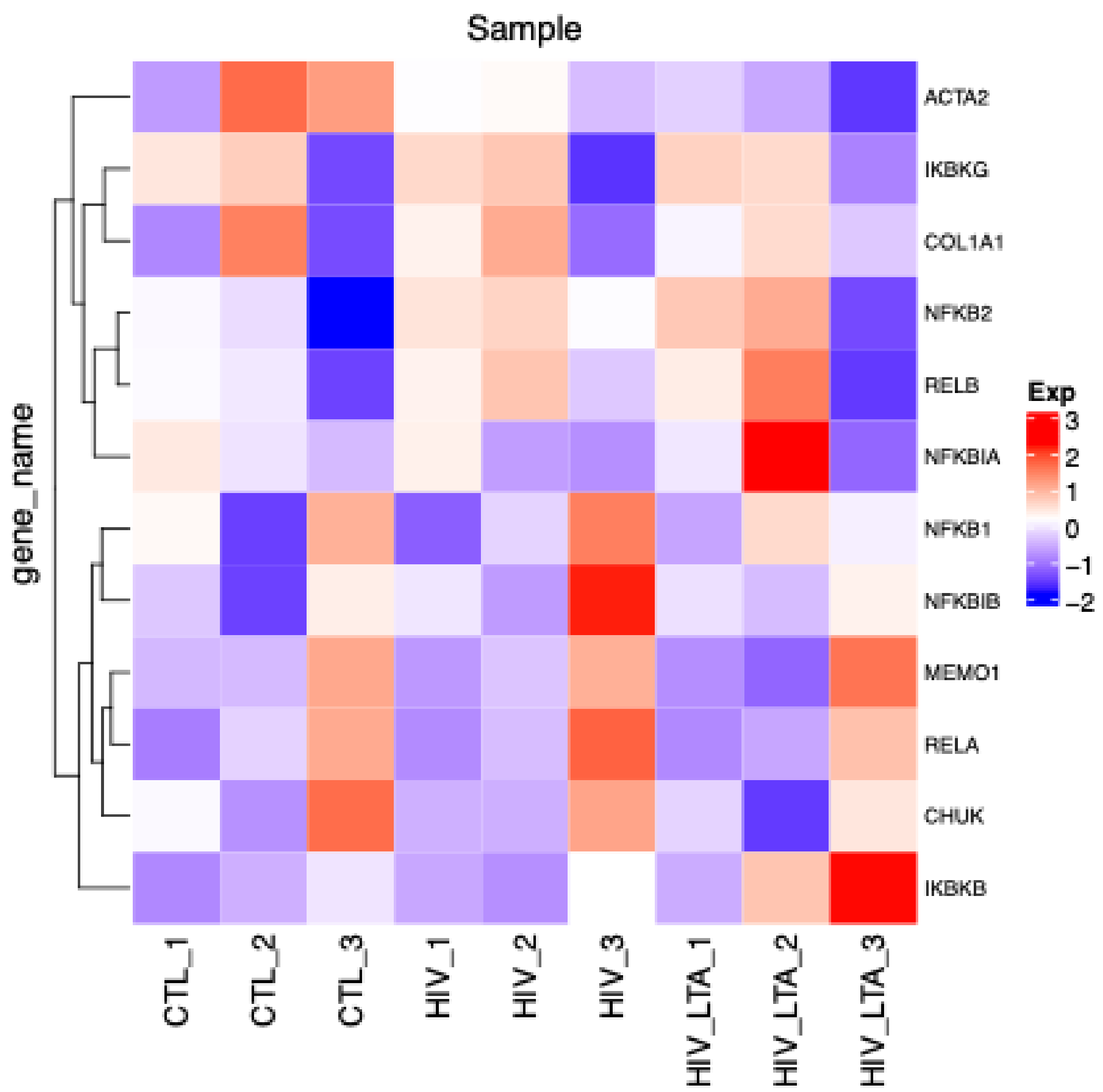


Heatmap analysis of NF-kB related genes across and major fibrogenic genes across the three conditions.
